# GWAS identifies 10 loci for objectively-measured physical activity and sleep with causal roles in cardiometabolic disease

**DOI:** 10.1101/261719

**Authors:** Aiden Doherty, Karl Smith-Byrne, Teresa Ferreira, Michael V. Holmes, Chris Holmes, Sara L. Pulit, Cecilia M. Lindgren

## Abstract

Physical activity and sleep disorders are established risk factors for many diseases, but their etiology is poorly understood, partly due to a reliance on self-reported evidence. Here we report a genome-wide association study (GWAS) of wearable-defined and machine-learned physical activity and sleep phenotypes in 91,112 UK Biobank participants, and self-reported physical activity in 351,154 UK Biobank participants. While the self-reported activity analysis resulted in no significant (p<5×10^−9^) loci, the analysis of objectively-measured traits identified 10 loci, 6 of which are novel. These 10 loci account for 0.05% of activity and 0.33% of sleep phenotype variation, but genome-wide estimates suggest that common variation accounts for ~12% of phenotypic variation, indicating high polygenicity. Heritability was higher in women than in men for overall activity (Δh^2^ = 4%, p=6.3×10^−5^), moderate intensity activity (6%, p=6.7×10^−8^), and walking (5%, p=2.6×10^−6^). Heritability partitioning, enrichment and pathway analyses all indicate the central nervous system plays a role in activity behaviours. Mendelian randomization in publicly available GWAS data and in 278,367 UK Biobank participants, who were not included in our discovery analyses, suggest that overall activity might be causally related to lowering body fat percentage (beta per SD higher overall activity: −0.44, SE=0.047, p=2.70×10^−21^) and systolic blood pressure (beta per SD: −0.71, SE=0.125, p=1.38×10^−8^). Our current results advocate the value of physical activity for the reduction of adiposity and blood pressure.

## INTRODUCTION

Physical inactivity is a global public health threat and is estimated to cost health-care systems ~$50 billion per year worldwide^1,2^. It is associated with a range of common diseases including multiple cardiometabolic outcomes such as obesity, type 2 diabetes, and cardiovascular diseases^3^. Alterations in sleep duration also associate with negative health outcomes, including cardio-metabolic diseases^4^ and psychiatric disorders^5^. In response to physical activity and sleep sharing a movement continuum, recent Canadian public health guidelines now recommend how youth should spend time in a variety of movement-related behaviours that make up the whole day^6^.

Twin and family studies have shown that self-reported daily physical activity is heritable, but with a large degree of heterogeneity in measurement methods and sample size (h^2^ range: 0%-78%)7. Emerging evidence shows a genetic contribution to physical activity, finding three variants^8^ associated with standard UK Biobank activity metrics^9^. Recent GWAS have also reported common variants that are associated with sleep duration^10–12^. However, GWAS of behavioural traits such as physical activity and sleep have mostly relied on self-reported data that are prone to measurement error and thus have limited statistical power. It is also possible that self-reported measures capture how well individuals perceive what they do, rather than what they actually do. Consequently, our understanding of the genes and biological pathways that underpin physical activity and sleep behaviours is still limited.

To better understand the genetic underpinnings of these behavioural traits, we performed a GWAS in 91,112 UK Biobank participants who were asked to wear activity trackers over a seven-day period. We used statistical machine learning to derive objective measures of physical activity and sleep from raw device data, which helped identify novel associations to genetic loci. The associated loci reveal pathways in the central nervous system that help advance our understanding of biological processes underlying human sleep and activity behaviours. Our study also reveals the importance of having objective measures at scale to detect genetic effects for behavioural traits. Mendelian randomization supports a role for physical activity in the reduction of body fat percentage and systolic blood pressure.

## RESULTS

### Physical activity and sleep traits in ~100,000 UK Biobank participants can be objectively measured through statistical machine learning of wearable sensor data

We examined data from 101,307 UK Biobank participants who agreed to wear a wrist-worn accelerometer for seven days^9^ and additionally underwent genome-wide genotyping and imputation^13^. As part of the UK Biobank Accelerometer Working Group, we generated continuous phenotypes representing overall activity and moderate-intensity activity time^9^ (**Methods**). Additionally, we applied a machine-learning model, using balanced random forests with Markov confusion matrices^14^, to identify which one of seven activity states an individual was in at any given time. We trained and evaluated the model in 57 free-living individuals to distinguish between activity states, with an overall classification score of kappa=0.79 in correctly predicting what activity state an individual is in for any given 30-second time period (**Table S1**). This is comparable to the level of performance (kappa=0.80) that we expect from humans defining reference activity labels in free-living evaluation datasets. We then decided to concentrate on sleep duration, sitting or standing, and walking behaviours as these occurred most commonly and are of most public health interest. We applied this model to over 100,000 UK Biobank participants^14^ (**Fig S1**).

### Novel loci associated with objective measures of physical activity, but not self-report

After quality control, 91,112 participants of European descent and 14,666,529 SNPs remained for subsequent genetic association analysis (**Methods**). Using similar quality control criteria, we also extracted information on self-reported physical activity (via the international physical activity questionnaire short-form^15^) for 351,154 UK Biobank participants of European descent (**Methods**). To maximise statistical power, we performed our analyses in BOLT-LMM^16^, which applies a linear mixed model to the data, allowing for inclusion of related individuals and those of varying genetic ancestry^17^. We additionally included assessment centre, genotyping array, age, and age squared as covariates.

Our analysis identified 10 significant loci (p<5×10^−9^) that were separated by at least 400 kilobases (kb) (**Table1, Fig S1**). We empirically derived this threshold (p<5×10^−9^) for genome-wide significance that considers multiple testing in densely imputed data^18^. Six loci were novel and these include: one for overall activity (rs59499656 near *SYT4*, p=2.0×10^−9^); two for sleep duration (rs7765476 near *LOC101928519*, p= 2.4×10^−13^; and rs2416963 near *MAPKAP1*, p= 6.9×10^−10^); two for sitting/standing (rs26579 near *MEF2C-AS2*, p=3.8×10^−11^; and rs113851554 near *MEIS1*, p= 1.4×10^−10^) none for walking; and one for moderate intensity activity (rs9938281 near *ZNF423*, p=1.8×10^−9^) (**Table 1, Fig S3**). The remaining associations were known: two for overall and moderate intensity activity^8^ (near *LINC02210-CRHR1* and *KANSL1-AS1*) and five for sleep duration^19–23,12^ (near *MEIS1*, *PAX8-AS1*, *MIR137*, *OLFM2,* and *MAPK8IP1P2*). Setting genome-wide significance at the conventional threshold of p<5×10^−8^ would have revealed 12 additional novel loci (**Table S2**). Of note, no loci were associated with self-reported physical activity status in our analysis (**Fig 1**). Nominal replication was seen for loci previously reported for overall and moderate intensity activity^8^ (near *DNAJB4* and *PML*), sleep^12^ (near *EBLN3P*, *PRKCB*, and *RPGRIP1L*), and sitting (near *LY6H*) (**Table S3**). No other findings were confirmed from previous GWAS that relied on self-report measures of physical activity and sleep (**Table S3**).

**Figure 1.**
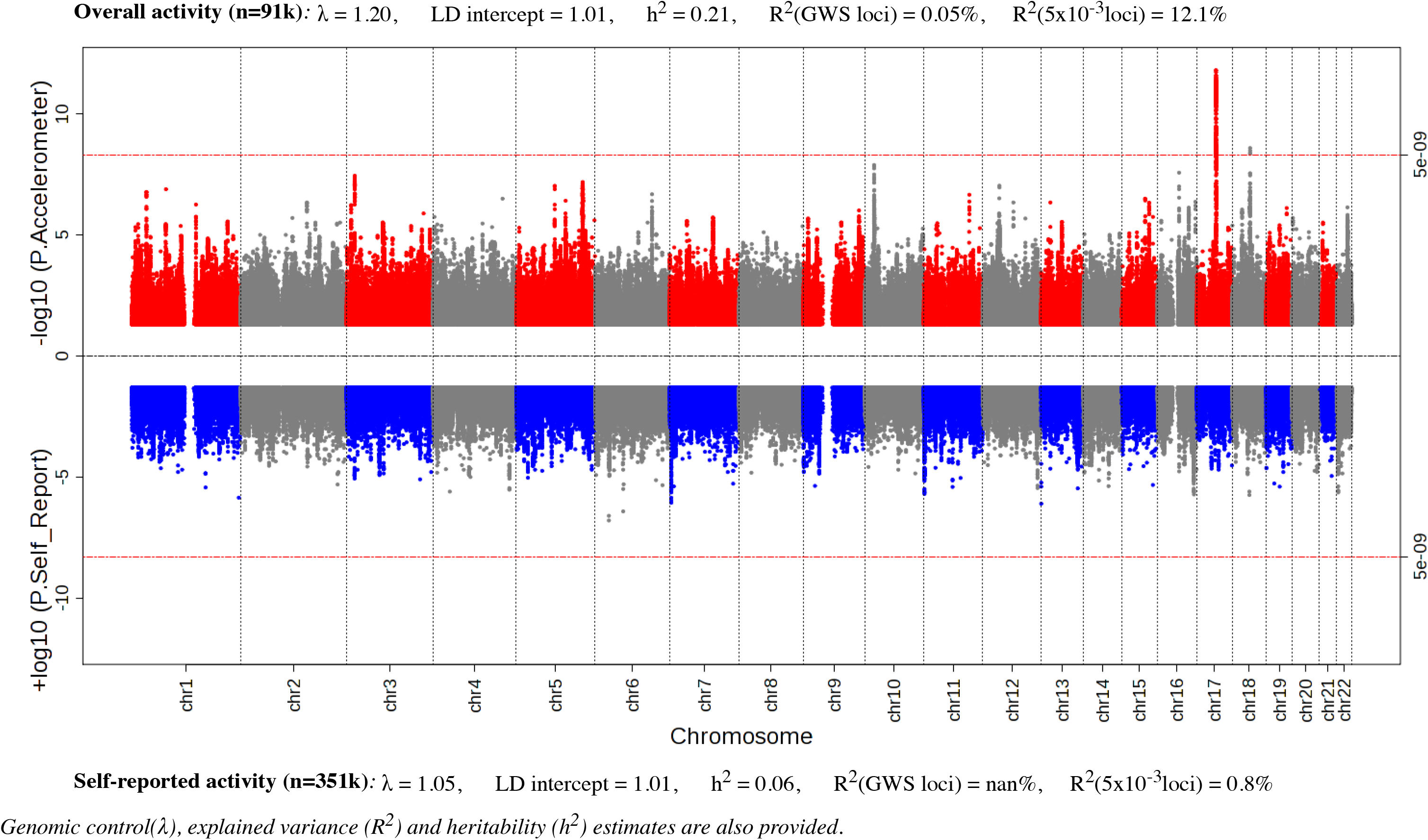
Miami plot of European sex-combined GWAS of physical activity in UK Biobank measured by accelerometer (*top, n=91,112*) and self-report (*below, n=351,154*).

**Table 1.** Genome-wide significant (5×10^−9^) loci associated with accelerometer-measured physical activity and sleep behaviours in 91,112 UK Biobank participants.

| Status | Trait | Locus | Chr | SNP | Position | Nearest gene | Allele (effect/other) | Effect allele frequency | beta | SE | P |
| --- | --- | --- | --- | --- | --- | --- | --- | --- | --- | --- | --- |
| Novel | Overall activity | 1 | 18 | rs59499656 | 40,768,309 | *SYT4* | A/T | 0.655 | -0.234 | 0.039 | $2.0 \times 10^{-09}$ |
| Novel | Sleep | 2 | 6 | rs7765476 | 19,069,450 | *LOC101928519* | T/G | 0.799 | 0.002 | 0.000 | $2.4 \times 10^{-13}$ |
| Novel | Sleep | 3 | 9 | rs2416963 | 128,241,414 | *MAPKAP1* | C/T | 0.589 | 0.002 | 0.000 | $6.9 \times 10^{-10}$ |
| Novel | Sit / stand | 4 | 5 | rs26579 | 87,985,295 | *MEF2C-AS2* | G/C | 0.415 | 0.002 | 0.000 | $3.8 \times 10^{-11}$ |
| Novel | Sit / stand | 5 | 2 | rs113851554 | 66,750,564 | *MEIS1* | G/T | 0.943 | -0.005 | 0.001 | $1.4 \times 10^{-10}$ |
| Novel | Moderate intensity activity | 6 | 16 | rs9938281 | 49,625,336 | *ZNF423* | A/G | 0.474 | -0.023 | 0.004 | $1.8 \times 10^{-09}$ |
| Known<sup>8</sup> | Overall activity | 7 | 17 | rs56194509 | 43,844,559 | *LINC02210-CRHR1* | T/G | 0.779 | -0.314 | 0.045 | $2.2 \times 10^{-12}$ |
| Known<sup>11</sup> | Sleep | 5 | 2 | rs113851554 | 66,750,564 | *MEIS1* | G/T | 0.943 | 0.006 | 0.001 | $2.6 \times 10^{-26}$ |
| Known<sup>11</sup> | Sleep | 8 | 2 | rs62158170 | 114,082,175 | *PAX8-AS1* | A/G | 0.783 | -0.003 | 0.000 | $1.6 \times 10^{-21}$ |
| Known<sup>Neale</sup> | Sleep | 9 | 1 | rs1198575 | 98,562,260 | *MIR137* | T/C | 0.190 | -0.002 | 0.000 | $8.3 \times 10^{-11}$ |
| Known<sup>21</sup> | Sleep | 10 | 19 | rs2303100 | 9,968,434 | *OLFM2* | C/T | 0.447 | -0.002 | 0.000 | $3.1 \times 10^{-10}$ |
| Known<sup>23</sup> | Sleep | 7 | 17 | rs7502280 | 43,670,221 | *MAPK8IP1P2* | T/G | 0.867 | 0.002 | 0.000 | $2.4 \times 10^{-09}$ |
| Known<sup>8</sup> | Moderate intensity activity | 7 | 17 | rs2957316 | 44,331,214 | *KANSL1-AS1* | T/C | 0.795 | -0.033 | 0.005 | $1.3 \times 10^{-11}$ |

Genomic control lambda across the GWAS ranged from 1.1 – 1.25, indicating modest deviation in the test statistics compared with the expectation. Estimation of the LD Score intercepts, using LD score regression^24^, revealed intercepts near 1.00 (range: 0.993 –1.015), indicating that the genomic inflation was due to a polygenic architecture rather than inflation due to uncorrected population structure. We performed approximate conditional analysis in each locus reaching genome-wide significance using Genome-wide Complex Trait Analysis^25^ (GCTA) but did not identify secondary signals in any locus. Gene-based analysis using FUMA^26^, with input SNPs mapped by position to 18,232 protein-coding genes, did not find additional loci to those already identified in the SNP association analysis (**Table S4**).

### Genetic architecture of physical activity and sleep

Using heritability estimates from BOLT-LMM^16^ (**Methods**), we found all traits to have modest heritability, ranging from 11% for walking to 22% for moderate intensity activity (**Fig S2**). Genome-wide significant loci account for 0.03% (sitting) to 0.33% (sleep) of phenotypic variance across the traits (**Fig S2**). Additional analyses testing for polygenic trait architecture indicate that SNPs with p-values well away from genome-wide significance add significantly to the phenotypic variance explained. For example, explained phenotypic variance ranged from 11.8% for walking to 13.2% for sleep when we selected independent SNPs (r^2^<0.1) with p<5×10^−3^ and distance >250kb from index SNPs.

We applied partitioned heritability^27^ analysis for tissue and functional categories using LD-score regression^24^. After taking the median value across all the 5 accelerometer-measured traits, we identified significant enrichments in the central nervous system and adrenal/pancreatic tissues (p<5×10^−3^, accounting for 10 cell types) (**Fig 2, S4**). Regions of the genome annotated as conserved across mammals showed the largest enrichment across functional categories and traits (p<2.38×10^−4^, accounting for 21 categories). These findings support a key role for physical activity and sleep behaviors throughout mammalian evolution.

**Figure 2.**
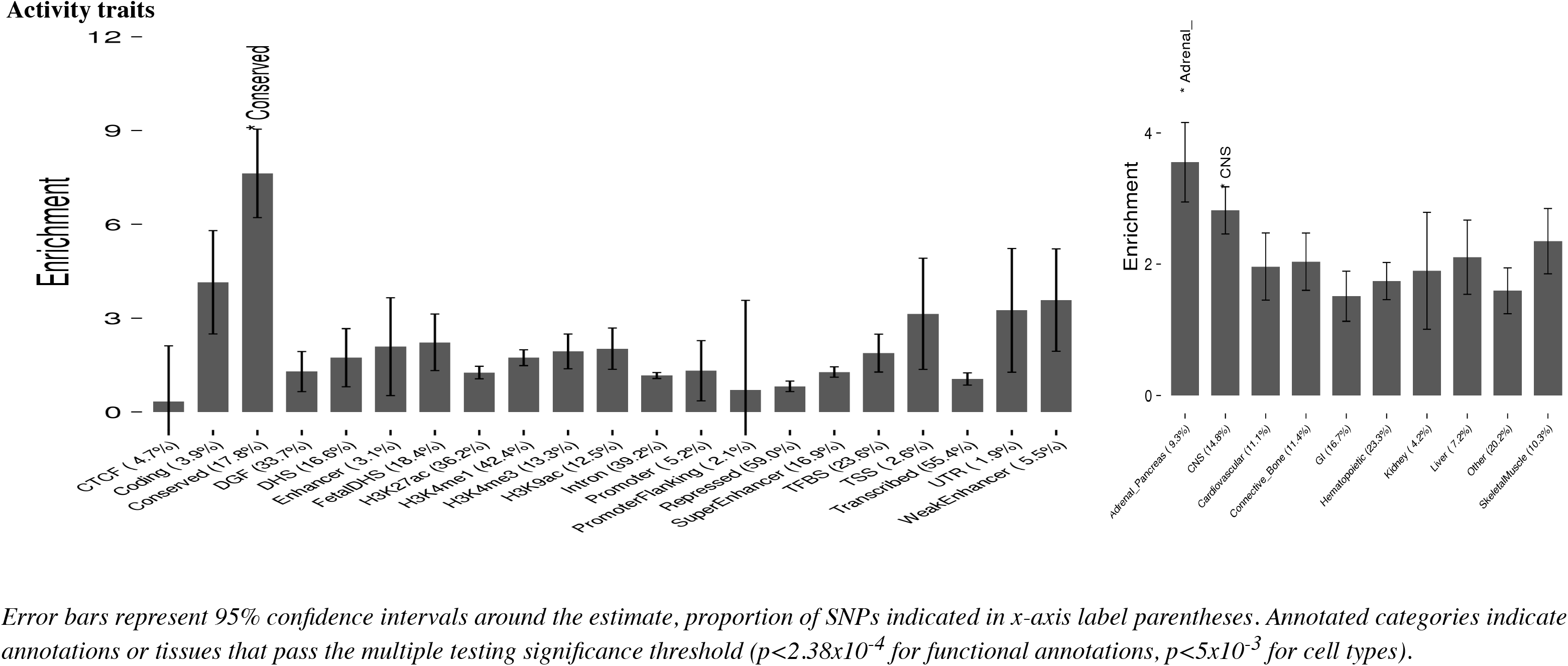
Heritability partitioning enrichment estimates across functional categories and tissues, for median value across 5 physical activity and sleep traits.

For all sleep and activity phenotypes, we used FINEMAP^28^ to identify credible sets of causal SNPs (meeting a log10 Bayes Factor > 2) in a 1Mb window around each index SNP, on the assumption that there is a maximum of 5 causal variants per locus. Four loci contained plausible causal variants in regions spanning <10kb (**Table S5**). One locus contained a single plausible causal variant near *MEIS1* in chromosome 2 for both sleep and sit/stand time.

### Sexual dimorphism in physical activity and sleep

Given that known sex differences exist for physical activity levels^9^ and muscle and fat mass distribution^29^, we investigated genetic sexual dimorphisms in our activity traits. Genetic correlations^30^ of trait architectures as measured using LD score regression^24^ showed strong correlation between men and women; correlations ranged from 0.89 for moderate activity to 0.96 for overall activity (**Fig S5**). In addition, there was no evidence of sexual dimorphism using strict criteria for heterogeneity of effects between men and women (p<5×10^−9^).

Heritability in women was higher than that in men for overall activity (Δh^2^ = 4%, p=6.3×10^−5^), moderate intensity activity (6%, p=6.7×10^−8^), and walking (5%, p=2.6×10^−6^) (**Fig S5**). We found no evidence for sexually-dimorphic heritability in sleep (p=0.12) and sit/stand behaviours (p=0.27). We found some evidence of sexually dimorphic variance explained in 4 of the 5 traits (**Fig S5**).

### Physical activity and sleep, and their relationship with other traits

While physical activity and sleep are established risk factors for multiple diseases from observational epidemiology, the extent to which their underlying genetic architectures are shared with disease phenotypes is unknown^7^. We conducted various analyses that included: 1) genome-wide genetic correlations between our traits and those from publicly-available GWAS data using LD score regression^24^ on the LD-Hub web resource^31^; 2) phenome-wide association (pheWAS) analysis for index SNPs using the Oxford Brain Imaging Genetics Server^32^ (big.stats.ox.ac.uk); and 3) a lookup of the NHGRI GWAS catalog^33^ for related SNPs associated with other diseases and traits (within 400kb and r^2^>0.2). Activity and sleep traits showed genetic associations with many other independent traits in LD score regression (n=158), pheWAS (n=115), and GWAS-catalog lookup (n=5) analyses (**Tables S6-8, Fig S6**). Associations with anthropometric and cognitive health traits were particularly common (**Note S1**). In general, increases in overall activity and walking phenotypes were genetically correlated with improved health status, while increases in sleep duration were negatively correlated with fluid intelligence scores and health status. Increases in sitting/standing time were genetically correlated with increased fluid intelligence scores, but decreased health status (**Note S1, Table S6**).

To investigate whether activity and sleep might contribute causally to disease outcomes, we performed Mendelian randomization (MR) analysis in 278,367 UK Biobank participants who were not included in our discovery analysis, and other publicly available GWAS summary data from the MR-Base web platform^34,35^ (**Methods**). Rather than selecting all significant traits from the previous genetic correlation analyses, we decided to only focus on major diseases and risk factors. These included cardiovascular disease (CHD, stroke, heart failure), type 2 diabetes, Alzheimer′s, major cancer subtypes, blood pressure, and anthropometric traits (BMI and body fat %). For the MR analysis of each trait, we selected instrument variables using traditional criteria for genome wide significance (p<5×10^−8^) and then followed a number of steps to denote potential causality with disease outcomes (**Methods, Note S2**). These included: maximum likelihood estimates^36^, leave-one-out analyses, robustness against pleiotropy via median and mode estimators, MR-Egger, and bi-directional tests^37^. We found consistent evidence of inverse relationships for overall activity with body fat percentage (beta per SD higher overall activity: −0.44, SE=0.047, p=2.70×10^−21^) and systolic blood pressure (beta per SD: −0.71, SE=0.125, p=1.38×10^−8^) (**Tables S9-11**). While there was evidence of bi-directional relationships (which is plausible), directionality analyses found evidence in support of activity being proximal to body fat percentage and systolic blood pressures (**Tables S11-12**).

### Potential functional and biological mechanisms for physical activity and sleep

We anticipated the use of objective measures would not only identify activity and sleep associated variants, but also provide insight on potential functional and biological mechanisms for these traits. First, we used DEPICT^38^ at suggestive loci with P<1×10^−5^ to identify gene sets enriched for movement phenotype associations; and tissues and cell types in which genes from associated loci are highly expressed. This analysis did not yield any significant results (FDR<0.05). Secondly, we used the FUMA^26^ web platform to identify tissues and gene pathways enriched for genetic signals (**Methods**). Here, we found brain tissues to be enriched, in particular at the cerebellum (p=3.4×10^−5^), cerebellar hemisphere (p=6.1×10^−5^), frontal cortex (p=9.1×10^−5^), brain cortex (p=1.5×10^−4^), and anterior cingulate cortex (p=3.9×10^−4^) (**Fig 3, S7**). We also identified pathways enriched in neurological disease, brain structure, and cognitive function, among other traits (**Fig S8**).

**Figure 3.**
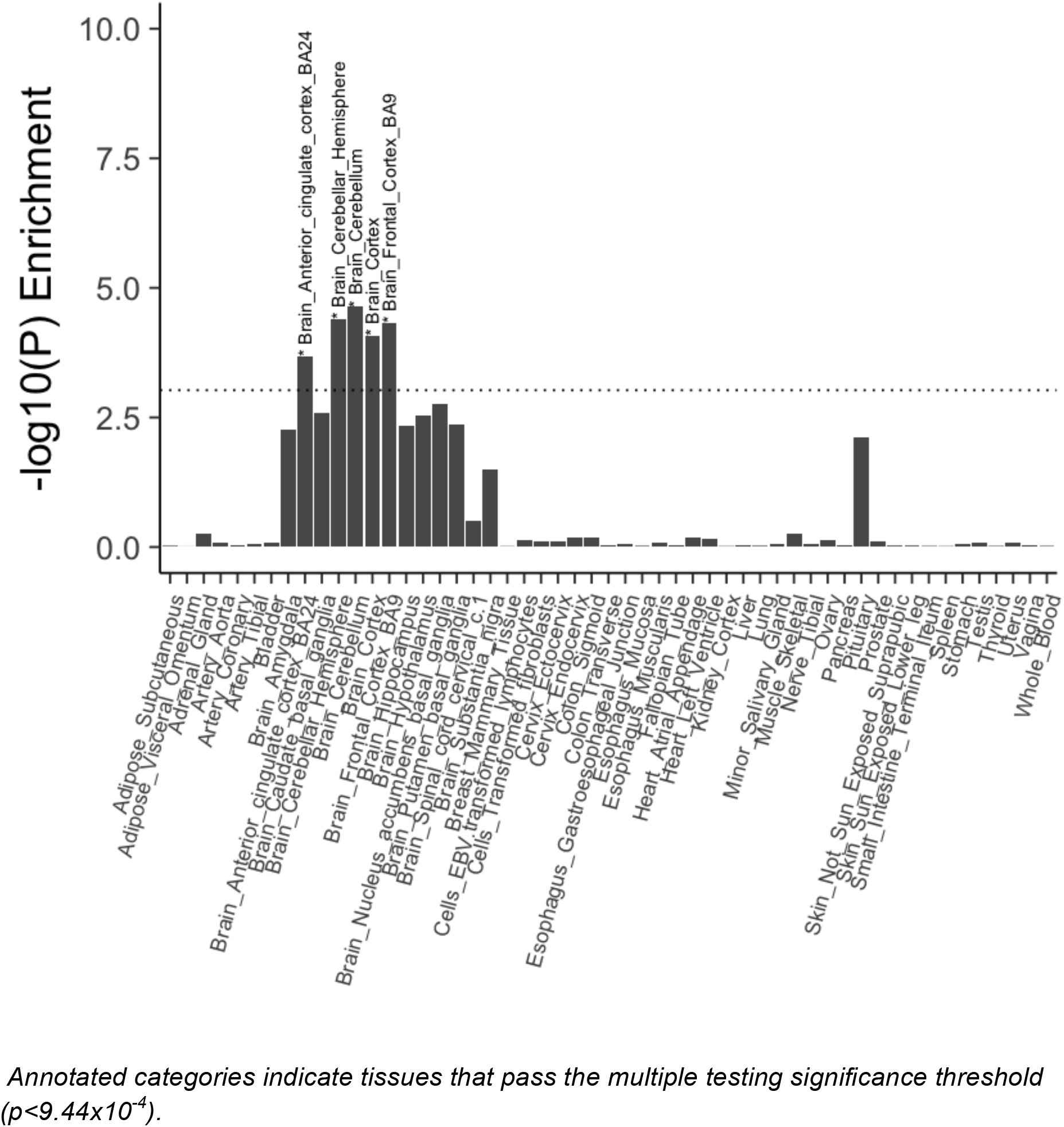
Tissue enrichment analysis using eQTL data from GTEx for median value across 5 physical activity and sleep traits.

## DISCUSSION

Our analysis of 91,112 individuals identified 10 loci significantly associated with objectively measured physical activity and sleep traits. While these loci alone account for less than 0.5% of the variation in physical activity, our analyses suggest that as much as 12% of activity variation might be accounted for by common (minor allele frequency > 1%) genetic variation. We found heritability was higher in women than in men for activity traits, but no differences were found for sleep and sitting/standing. As anticipated, physical activity and sleep have polygenic architectures and share biology with multiple traits including intelligence, education, obesity, and cardiometabolic diseases.

Our results advocate the value of physical activity for the reduction of adiposity and blood pressure. Until now, it has been difficult to discern whether associations between activity and disease are truly causal or biased due to reverse causality, confounding, and self-report measurement error associated with traditional observational studies^39^. Ascertainment bias is also an issue in randomised controlled trials of physical activity, due to difficulties in blinding participants to behavioral interventions^40^. Debate on whether obesity is a determinant of physical activity, or vice-versa, has previously lacked information due to a paucity of instrument variables for physical activity^7,41^. Our MR analyses suggest that higher levels of physical activity might causally lower body fat percentage and systolic blood pressure. We note that our current MR estimates are based on a limited number of SNPs. While this number of markers may be insufficient to comprehensively instrument a complex phenotype, early studies of body mass index used only one SNP and obtained reliable estimates^42^.

We observed an important role for the central nervous system with respect to physical activity and sleep, consistent with emerging literature^8^. In addition to these findings from heritability partitioning, enrichment and pathway analyses; our gene-set and pheWAS analyses showed genetic overlap with neurodegenerative diseases, mental health wellbeing, brain structure, function, and connectivity. Intriguingly, a recent GWAS of BMI highlighted a key role for neurological pathways in overall fat distribution^43^ while a GWAS of fat distribution as measured by waist-to-hip ratio highlight adipose pathways^44^. It is therefore possible that our findings might be driven through obesity, as we observed strong genetic correlations between activity and obesity traits. However, as this is one of the first large-scale physical activity genetic studies, we have likely only begun to understand the complex interplay between activity, cardiometabolic phenotypes, and neurologic health.

Our work demonstrates the importance of large-scale objective measures to identify genetic associations with behavioural phenotypes. For example, we found no loci were associated with self-reported activity despite the much larger sample of 351,154 UK Biobank participants. Our heritability estimates for sleep were consistent with previous self-reported literature^19–21^, but we found that self-reported data significantly underestimated heritability for physical activity^8^ (~20% vs ~5%). While large-scale self-reported measures have appeared robust for other phenotypes such as birthweight^45^ and major depression status^46^, it appears that measurement error is a considerable obstacle when assessing behavioural traits. Recall, comprehension, and social-desirability bias are more prevalent in behavioural traits such as physical activity, where phenotypic agreement is generally poor between subjective and objective measures (typically r<0.2)^47^.

This study has many strengths including: objectively measured behavioural phenotypes validated in free-living environments, use of linear mixed models, and large sample sizes, all key to maximise power in genetic discovery analyses; and Mendelian Randomization analyses, for delineating causal relationships. However, given the cohort age range (45-80 years) and inclusion of European ancestry individuals only, we cannot necessarily assume our results generalise to other age groups or ancestral populations. Larger studies of common genetic variation, and studies in non-European populations, will help in replicating these findings, as well as continue to further refine the biology of physical activity and the diseases it might cause. To account for the likely U-shape relationship between sleep and disease outcomes^48^, rather than a linear relationship as assumed in current models, future work will be also required for MR sleep analysis.

In summary, we conducted the largest GWAS of wearable-defined and machine-learned sleep and physical activity phenotypes to date, and identified 10 trait-associated loci. Our analysis shows shared genetic pathways with multiple traits including intelligence-related phenotypes and cardiometabolic disease. In support of national and international clinical guidance, this study provides strong evidence for physical activity in the prevention of adiposity and blood pressure.

## METHODS

### Participants

Study participants were from the UK Biobank^13^ where a subset of 103,712 participants agreed to wear a wrist-worn accelerometer for a seven day period between 2013-2015^9^. This study (UK Biobank project #9126) was covered by the general ethical approval for UK Biobank studies from the NHS National Research Ethics Service on 17th June 2011 (Ref 11/NW/0382). For the development and free-living evaluation of accelerometer machine learning methods, we used a validation set of 57 participants recruited to the CAPTURE-24 study where adults aged 18-91 were recruited from the Oxford region in 2014-2015^14^. Participants were asked to wear a wrist-worn accelerometer (same as in UK Biobank) for a 24-hour period and then given a £20 voucher for taking part in this study that received ethical approval from University of Oxford (Inter-Divisional Research Ethics Committee (IDREC) reference number: SSD/CUREC1A/13-262).

### Device

Participants wore an Axivity AX3 wrist-worn triaxial accelerometer on their dominant hand at all times over a seven-day period. It was set to capture tri-axial acceleration data at 100Hz with a dynamic range of +-8g. This device has demonstrated equivalent output^49^ to the GENEActiv accelerometer which has been validated using both standard laboratory and free-living energy expenditure assessment methods^50,51^. For data pre-processing we followed procedures that we developed as part of the UK Biobank accelerometer data processing expert group^9^, that included device calibration, resampling to 100Hz, and removal of noise and gravity.

### Development and validation of machine-learned phenotypes

To create a “ground truth” to extract sleep, sitting/standing, and walking time from sensor data, participants in the validation study also wore cameras in natural free-living, rather than constrained laboratory, environments. Wearable cameras automatically take photographs every ~20 seconds, have up to 16 hours battery life and storage capacity for over one week’s worth of images. When worn, the camera is reasonably close to the wearer’s eye line and has a wide-angle lens to capture everything within the wearer’s view. Each image is time-stamped so duration of sitting/standing, waking, and a range of other physical activity behaviours can be captured^52^. To extract sleep information, participants were asked to complete a simple sleep diary, as used in the Whitehall study, which consisted of two questions^53^: ‘*what time did you first fall asleep last night?*’ and ‘*what time did you wake up today (eyes open, ready to get up)?*’. Participants were also asked to complete a Harmonised European Time Use Survey diary^54^, and sleep information from here was extracted in cases where data was missing from the simple sleep diary.

For every non-overlapping 30-second time window, we then extracted a 126-dimensional feature vector representing a range of time and frequency domain features^14^. For activity classification we used random forests^55^ which offer a powerful nonparametric discriminative method for classification that offers state-of-the-art performance^56^. Random forests are able to classify data points, but do not have an understanding of our data as having come from a time series. Therefore we then used a hidden Markov model^57^ (HMM) to smooth our predictions. This smoothing corrects erroneous predictions from the random forest, such as where the error is a blip of one activity surrounded by another and the transitions between those two classes of activity are rare. This allowed us to train a model using all free-living ground truth data, achieving 86% accuracy across classes of interest^14^ (**Table S1**).

### Sleep and physical activity phenotypes in UK Biobank

To predict **sleep, sitting/standing, and walking**, we applied our machine learning method to predict behaviour for each 30-second epoch in 103,712 UK Biobank participants’ accelerometer data. For any given time window (e.g., one hour, one day, etc.) the probability of a participant engaging in a specific behaviour type was expressed as the number-of-epoch-predictions-for-class divided by the number-of-epochs. For **overall activity** levels, we selected average vector magnitude for each 30-second epoch, which is the recommended variable for activity analysis^9^. This variable has been shown to account for 44-47% explained variance versus combined sensing heart-rate + trunk-acceleration (a proxy for free-living physical activity energy expenditure) in 1695 UK adults^51^. To infer time spent in **moderate intensity activity**, we considered all epoch periods at or greater than 100 milli-gravity units. This threshold has been shown to be equivalent to 3 METs in laboratory experiments^58^.

Device non-wear time was automatically identified as consecutive stationary episodes lasting for at least 60 minutes^9^. These non-wear segments of data were imputed with the average of similar time-of-day data points, for each behaviour prediction, from different days of the measurement. We excluded participants whose data could not be calibrated (n=0), had too many clipped values (n=3), had unrealistically high values (average vector magnitude > 100mg) (n=32), or who had poor wear-time (n=6,860). We defined minimum wear time criteria as having at least three days (72 hours) of data and also data in each one-hour period of the 24-hour cycle^9^.

We also extracted information on self-reported physical activity (via the international physical questionnaire short-form^15^). Here, self-reported walking (field #874), moderate (#894), and vigorous (#914) activity duration were converted into MET-hours (a unit of physical activity energy expenditure) following procedures described elsewhere^15^. Rather than ignoring participants at the bottom end of the self-reported activity spectrum^8^, we included such participants who answered ‘0’ to number of days in a week that they walked (field #864) or did moderate (#field 884) or vigorous (field #904) intensity activity (for at least 10 minutes). Participants were removed from analysis if they answered “Do not know” or “prefer not to answer” to the aforementioned questions (n=117,055).

### Genotyping and quality control

The UK Biobank provides ~92 million variants, including imputation based on UK10K haplotype, 1000 Genomes Phase 3, and Haplotype Reference Consortium (HRC) reference panels^59^. We excluded SNPs not in the HRC panel (~53M), with MAF < 0.1% (~24M), and with an imputation R^2^<0.3 (~14k). For self-reported physical activity, we removed participants who self-reported as being non-white (n=18,671), had abnormal genetic versus self-reported sex mismatches (n=146) or sex chromosome aneuploidy (n=383). This left 351,154 UK Biobank participants of European descent for subsequent genetic association analysis. For accelerometer measured traits, we removed participants who self-reported as being non-white (n=3,192), had abnormal genetic versus self-reported sex mismatches (n=36) or sex chromosome aneuploidy (n=71). This left 91,112 participants of European descent (39,972 men; 51,140 women), and 14,666,529 SNPs, for subsequent genetic association analysis.

### Identification of loci associated with activity and sleep traits

To improve our power to detect associations, we used BOLT-LMM^16^ to perform linear mixed models analysis. As this adjusts for population structure and relatedness, we could include many additional individuals (13,320 for accelerometer analysis and ~70,000 for self-report analysis) than if following the common practice of analysing a reduced set of unrelated white British individuals using linear regression^60^. We included assessment centre, genotyping array, age, and age squared as covariates. Sensitivity analysis showed we did not need to include principal components of ancestry as covariates in our linear mixed models. We then used PLINK^61^ (version 1.9) to exclude candidate variants (p<1×10^−5^) that deviated from Hardy-Weinberg equilibrium (p<1×10^−7^). To identify genome-wide-significant (GWS) loci, we defined a distance criterion of +-400kb surrounding each GWS peak (p<5×10^−9^). We empirically derived this threshold for genome-wide significance that considers multiple testing and densely imputed data^18^. For reporting, we also identified loci using a traditional GWS value of P<5×10^−8^.

To identify novel loci, we considered index SNPs falling outside 400kb of a SNP previously associated with one of the following: self-report sleep duration or physical activity from the NHGRI-EBI GWAS catalog^33^; accelerometer measured overall and moderate intensity activity loci reported in a bioRxiv paper that used 67,000 UK Biobank participants^8^; accelerometer-measured sleep duration genes tested for replication in a bioRxiv paper that used 83,726 UK Biobank participants^23^; self-reported overall activity loci reported in bioRxiv papers using ~277,000 and ~120,000 UK Biobank participants respectively^8,62^; self-reported sleep duration GWAS results in a bioRxiv paper that used 384,317 UK Biobank participants^12^; and sleep duration (field #1160), television watching time as a proxy for sitting (field #1070), and walking (field #874) GWAS results in ~330,000 UK Biobank participants published by the Neale lab at the Broad Institute.

To identify additional signals in regions of association, we performed approximate joint and conditional SNP association analysis in each locus using the Genome-wide Complex Trait Analysis^25^ (GCTA) tool. Any lead SNPs identified in a known high-LD area^63^ between 43.5-45.5mb at chromosome 17 were treated as a single large locus in GCTA analysis.

Gene based analysis were performed with MAGMA^64^ v1.6, as implemented on the FUMA^26^ web platform. Input SNPs were mapped by position to genes obtained from 18,232 protein-coding genes obtained via Ensembl^65^ build 85. Bonferroni-correction was used to define genome-wide significance. To identify novel loci, we looked for genes more than 400kb away from loci identified in the above SNP association analysis.

### Genetic architecture of activity and sleep traits

To estimate heritability of each trait, we used BOLT-LMM^16^ for computational efficiency, after sensitivity analysis showed negligible differences in estimates between BOLT-LMM and restricted maximum likelihood analysis^66^ implemented in BOLT-REML. We applied partitioned heritability^27^ analysis across the phenotypes by tissue and functional category using LD-score regression^24^ with the LDSC tool. Significant enrichments for individual traits were identified using thresholds of p<1×10^−3^ for tissues (i.e. p<0.05/10 cell types/5 traits) and p<3.57×10^−4^ for functional categories (i.e. p<0.05/28 categories/5 traits). We also investigated enrichments in the median value across all accelerometer-measured traits.

To calculate the explained phenotypic variance for each trait, we used the PRSice^67^ tool to generate polygenic risk scores for the lead SNP in GWS loci and also for all SNPs with p<5×10^−3^, distance > 250kb from index SNPs, and r^2^< 0.1. The same participant inclusion criteria were used as for association analysis.

To identify plausible causal SNPs associated with sleep and activity phenotypes, we used the FINEMAP^28^ software. Configurations of plausible causal SNPs from a 1Mb window around each genome-wide significant locus were calculated on the assumption that there were a maximum of five causal variants per locus. Across all loci, we defined plausible causal SNPs as those meeting a log10 Bayes Factor > 2.

### Sexual dimorphism in activity and sleep traits

To investigate potential sources of sex heterogeneity, we ran the aforementioned genome wide association analyses for men and women independently. Genetic correlations^30^ of trait architectures between men and women were measured using LD score regression^24^ for each phenotype. We also tested for heterogeneity of effect estimates^29^ between men and women for all SNPs using the EasyStrata^68^ tool, with P<5×10^−9^ to assess significance. Heritability differences between men and women for each trait were assessed by extracting a two-tailed p-value from the following z-score (where ‘var’ indicates variance):

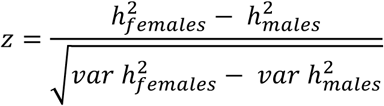

To estimate genetic contributions to phenotypic variance for men and women separately, we selected index SNPs from loci with a less stringent p<5×10^−7^, due to smaller sample sizes within strata.

### Associations of activity and sleep traits with other traits

To estimate genetic correlations between our movement phenotypes and other complex traits and diseases, we used LD score regression^24^ implemented in the LD-Hub web resource^31^. We did not test moderate intensity activity as it had a strong genetic correlation with overall activity (rg = 0.97, se = 0.004, **Fig S9**). To assess significance, we corrected for 3,328 tests (activity/sleep/sitting/walking x 832 phenotypes available on LD-Hub) with p<1.5×10^−5^.

To examine whether the genome-wide significant SNPs identified in our analysis affected other traits, we used the Oxford Brain Imaging Genetics Server (big.stats.ox.ac.uk^32^) to perform a phenome-wide association study (PheWAS) on almost 4,000 traits in UK Biobank participants. This included GWAS results of ~2,000 phenotypes in ~330,000 UK Biobank participants published by the Neale lab at the Broad Institute, and the remaining traits were brain imaging-derived phenotypes measured on a subset of ~10,000 UK Biobank participants^32^. To assess significance, we corrected for 40,000 tests (10 loci x 4,000 traits) with p<1.25×10^−6^. Additionally, we extracted previously reported GWAS associations within 400kb and r^2^>0.2 of accelerometer index SNPs from the NHGRI GWAS catalog^33^.

To investigate whether activity and sleep might causally contribute to disease outcomes, we performed Mendelian Randomization (MR) analysis. Rather than selecting all significant traits identified in other correlation analysis, we decided to concentrate on major diseases and peripheral risk factors. These included vascular disease (CHD, stroke, heart failure), diabetes, Alzheimer′s, major cancer subtypes, blood pressure, and anthropometric traits (BMI and body fat %). Disease phenotypes were prepared following similar procedures as used for UK Biobank variables in the LD-Hub web resource^31^. We defined hypertensive cases as individuals with systolic blood pressure of >140 mmHg, or a diastolic blood pressure of >90 mmHg, or the report of blood pressure medication usage. For the analysis of systolic and diastolic blood pressure, we corrected blood pressure measures in people on antihypertensive drugs by adding 15 mmHg to systolic and 10 mmHg to diastolic blood pressure, in keeping with the approach taken by genome-wide association studies^69^. Similar to the LD-Hub web resource^31^, we used linear regression analyses with sex and the first 10 principal components as covariates. For linear regression, we used the bgenie tool^59^. We removed participants from the accelerometer discovery sample (n=91,112), those who self-reported as being non-white British (n=68,428), had abnormal genetic versus self-reported sex mismatches (n=192) or sex chromosome aneuploidy (n=652). We also removed 48,658 participants due to relatedness. This left 278,367 UK Biobank participants who were not included in our discovery analysis, and other publicly available GWAS summary data from the MR-Base web platform^34^. Where possible, estimates were meta-analysed using a fixed effects model (inverse variance weighted average).

For analysis we retained index SNPs with p<5×10^−8^ that were pruned for LD (r^2^ < 0.001) and more than 10,000 kb apart. We then followed a number of steps to denote potential causality with disease outcomes (**Note S2**). We used the maximum likelihood-based approach as our primary source of MR estimates. This is based on published simulation results suggesting that causal estimates obtained from summarized data using a likelihood-based model are almost as precise as those obtained from individual-level data^36^. Only likelihood-based risk estimates that were significant after correction using a false-discovery rate of <5% were considered. The potential effect of pleiotropy was evaluated by three complementary approaches, namely weighted median and weighted mode estimation^70,71^, and the regression intercept from the MR-Egger method^72^. The sensitivity of causal inference to any individual genetic variant was tested by leave-one-out analysis. The Steiger test was used to provide evidence for the causal direction of the effect estimates^73^. Additionally, for MR associations that appeared robust to sensitivity and pleiotropy analyses, bidirectional MR was conducted to assess the direction of the causal estimate. All MR analyses were conducted in R using the TwoSampleMR package^34^.

### Investigating functional and biological mechanisms for activity and sleep traits

To investigate potential biological mechanisms underlying physical activity and sleep, we used DEPICT^38^ at suggestive loci with p<1×10^−5^ to identify: the most likely causal gene; reconstituted gene sets enriched for movement phenotype associations; and tissues and cell types in which genes from associated loci are highly expressed. Next, we used the FUMA web platform^26^ to perform tissue enrichment analysis where the full distribution of SNPs was tested with 53 specific tissue types, based on GTEx^74^ data. To identify significant enrichments, we accounted for multiple testing across 53 tissues and 5 traits (p<1.88×10^−4^). To then identify pathways implicated by the activity and sleep associated loci, we used FUMA to perform hypergeometric tests on genes from these loci to investigate over-representations in genes predefined from the GWAS-catalog.

## Data and code availability

The summary phenotype variables that we have constructed will be made available as a part of the UK Biobank dataset at http://biobank.ctsu.ox.ac.uk/crystal/label.cgi?id=1008. All data processing, feature extraction, and machine learning, code will be available at https://github.com/activityMonitoring.

## ACKNOWLEDGEMENTS

This research has been conducted using the UK Biobank resource under application number 11867. We would like to thank all participants for agreeing to volunteer in this research. Tugce Karaderi and Sven Hollowell contributed to data preparation. Matthew Willetts and Louis Aslett contributed to the development of machine learning methods to identify activity phenotypes. We would also like to thank Benjamin Neale at the Broad Institute for advice on statistical testing for heritability differences between stratified groups. For data collection at Oxford and annotation we would like to acknowledge the support of Jonathan Gershuny, the UK Economic and Social Research Council (grant number ES/L011662/1), Charlie Foster, Paul Kelly, Teresa Harms, Emma Thomas, Karen Milton, and Wong Tsz Yan. The UK Biobank Activity Project and the collection of activity data from participants was funded by the Wellcome Trust (https://wellcome.ac.uk/) and the Medical Research Council (http://www.mrc.ac.uk/). The analysis was supported by: the NIHR Biomedical Research Centre, Oxford [AD, CML]; the British Heart Foundation Centre of Research Excellence at Oxford (http://www.cardioscience.ox.ac.uk/bhf-centre-of-research-excellence) [grant number RE/13/1/30181 to AD]; the Li Ka Shing Foundation (http://www.lksf.org/) [to AD, CML]; Welcome Trust-SSI/John Fell funds Oxford [CML]; Widenlife [CML]; and NIH [grant number CRR00070 CR00.01 to CML]. We would also like to acknowledge the use of the University of Oxford Advanced Research Computing (ARC) facility in carrying out this work. http://dx.doi.org/10.5281/zenodo.22558 The MRC and Wellcome Trust played a key role in the decision to establish UK Biobank, and the accelerometer data collection. No funding bodies had any role in the analysis, decision to publish, or preparation of the manuscript.

## AUTHOR CONTRIBUTIONS

AD, CH, and CML conceived this study. AD developed and implemented the analyses, with input from TF, and SP. KS-B developed and implemented the Mendelian Randomization analysis with input from MH. AD wrote the paper. AD, CML, SP, MH, CH, TF, and KS-B were involved in interpreting results and editing the manuscript.

